# KinOrtho: a method for mapping human kinase orthologs across the tree of life and illuminating understudied kinases

**DOI:** 10.1101/2021.03.05.434161

**Authors:** Liang-Chin Huang, Rahil Taujale, Nathan Gravel, Aarya Venkat, Wayland Yeung, Dominic P Byrne, Patrick A Eyers, Natarajan Kannan

## Abstract

Protein kinases are among the largest druggable family of signaling proteins, involved in various human diseases, including cancers and neurodegenerative disorders. Despite their clinical relevance, nearly 30% of the 545 human protein kinases remain highly understudied. Comparative genomics is a powerful approach for predicting and investigating the functions of understudied kinases. However, an incomplete knowledge of kinase orthologs across fully sequenced kinomes severely limits the application of comparative approaches for illuminating understudied kinases. Here, we propose KinOrtho, a query- and graph-based orthology inference method that combines full-length and domain-based approaches to map one-to-one kinase orthologs across 17 thousand species. Using multiple metrics, we show that KinOrtho performed better than existing methods in identifying kinase orthologs across evolutionarily divergent species and eliminated potential false positives by flagging sequences without a proper kinase domain for further evaluation. We demonstrate the advantage of using domain-based approaches for identifying domain fusion events, highlighting a case between an understudied serine/threonine kinase TAOK1 and a metabolic kinase PIK3C2A with high co-expression in human cells. We also identify evolutionary fission events involving the understudied OBSCN kinase domains, further highlighting the value of domain-based orthology inference approaches. Using KinOrtho-defined orthologs, Gene Ontology annotations, and machine learning, we propose putative biological functions of several understudied kinases, including the role of TP53RK in cell cycle checkpoint(s), the involvement of TSSK3 and TSSK6 in acrosomal vesicle localization, and potential functions for the ULK4 pseudokinase in neuronal development.

The well-curated kinome ortholog set can serve as a valuable resource for illuminating understudied kinases, and the KinOrtho framework can be extended to any gene-family of interest.

## Introduction

Since the completion of the human genome project, thousands of species have been fully sequenced (Adams et al. 1991), providing a broader coverage of species diversity across the tree of life. “Moonshot” approaches, such as the Earth BioGenome Project (EBP), aim to catalog, and then characterize, genomes across eukaryotic biodiversity during the next decade (Lewin et al. 2018). The acquisition of genomic (and their associated proteomic) datasets enables the accurate prediction of protein functions through ever-deeper evolutionary analysis of related sequences (Weiss 1993). Protein kinases transfer the gamma phosphate group from ATP to an expanding subset of amino acids in their regulatory targets (Hardman et al. 2019). They can be distinguished from other mechanistically related enzymes, such as metabolic and glycan-modifying kinases (Shrestha et al. 2020b). Protein kinases represent one of the largest druggable families of signaling proteins that are abnormally regulated in various human diseases, including most human cancers (Hopkins and Groom 2002; Arslan et al. 2006). The human genome encodes nearly 550 protein kinase-related genes (collectively referred to as the human kinome) that have been broadly classified into major groups and families (Manning et al. 2002; Wilson et al. 2018). A majority of the human kinome members have been functionally characterized in multiple model organisms; however, nearly 30% of human kinases remain understudied, despite multi-organism knowledge of their primary sequence. These orphan kinases are collectively referred to as “dark” kinases (Health 2019) (Jensen PubMed score (Nguyen et al. 2017) < 50, no R01s, and PubTator score (Wei et al. 2013) < 150), and many contain clear orthologs in a majority of eukaryotic genomes, suggesting essential biological functions across life. A major focus of the Illuminating the Druggable Genome (IDG; https://commonfund.nih.gov/idg/index) consortium is to characterize the functions of these understudied proteins as a conceptual starting point for developing new drugs for a wide range of diseases such as cancer, neurodegenerative and auto-immune disorders that are associated with abnormal kinome signaling (Moret et al. 2020).

Comparative genomics is a powerful approach for inferring gene functions and is based on the assumption that genes descended from the same ancestor are likely to retain commonly shared functions (Fitch 1970; Wei et al. 2002). These gene descendants are called orthologs and paralogs, two major types of homologs related to speciation and duplication events, respectively (Fitch 2000). Paralogs can be further defined as in-paralogs and out-paralogs: the former arises from duplication after speciation, while the latter arises from duplication before speciation (Koonin 2005). The concept of “one-to-one” orthologous relationships (one protein in one species versus one protein in the other species) has been extended to “one-to-many” or “many-to-many” relationships and are collectively termed orthologous groups (Tatusov et al. 1997). Co-orthologs are defined as a pair of genes from the same orthologous group but different species (O’Brien et al. 2005). Given the importance of these relationships for functional analysis, several orthology inference methods have been developed. We have previously used these approaches to analyze canonical protein kinases and pseudokinases, including a broad survey of pseudoenzymes (Ribeiro et al. 2019), pseudokinases (Kwon et al. 2019), and a variety of understudied kinases whose biological function remains unknown, despite conservation in various eukaryotic lineages (Bailey et al. 2015; Eyers et al. 2017; Shrestha et al. 2020a).

Current orthology inference methods can be broadly classified into two major categories: tree-based methods (Vilella et al. 2009; Kaduk and Sonnhammer 2017; Huerta-Cepas et al. 2019; Mi et al. 2019) and graph-based methods (Li et al. 2003; Lechner et al. 2011; Altenhoff et al. 2013; Sonnhammer and Ostlund 2015; Train et al. 2017; Cosentino and Iwasaki 2019; Emms and Kelly 2019; Nevers et al. 2019; Derelle et al. 2020). Tree-based methods, such as EnsemblCompara (Vilella et al. 2009), construct reconciled trees based on gene trees and corresponding species trees while graph-based methods identify hypothetical orthologs, expand the graph by adding in-paralogs and co-orthologs, and then cluster them into orthologous groups. Tree-based methods are generally more accurate than graph-based methods, depending on the accuracy of species trees (Gabaldon 2008; Trachana et al. 2011). However, tree-based methods are computationally expensive in terms of time, limiting the exploration of thousands of species across the tree of life (Gabaldon 2008). In contrast, graph-based methods such as OrthoMCL (Li et al. 2003) are faster, but the increased speed is usually achieved at the cost of reduced sensitivity.

Most orthology inference methods rely on time-consuming all-vs-all sequence similarity searches across full-length gene or protein sequences across entire genomes. As such, these methods are not designed for focused analysis on individual gene families. Within large protein families, such as the protein kinase superfamily, traditional orthology inference methods possess high false-positive rates since they do not consider the conservation of known functional domains. It can end up identifying sequences as putative orthologs that almost certainly lack the classical bilobal kinase domain. In contrast, domain-based methods, such as Hierarchical grouping of Orthologous and Paralogous Sequences (HOPS) (Storm and Sonnhammer 2003), FlowerPower (Krishnamurthy et al. 2007), Domain based Detection of Orthologs (DODO) (Chen et al. 2010), Microbial Genome Database (MBGD) (Uchiyama et al. 2015), and Domainoid (Persson et al. 2019), are tailored to identify evolutionary relationships based on functionally relevant regions, notably domains, of a protein. However, the performances of these methods are reliant on the annotation of domains based on prior knowledge, thereby making it challenging to identify novel domains and relationships. A good example of this is the discovery of atypical kinases with very low sequence identity compared to search sequences, such as the atypical SelO kinase (Sreelatha et al. 2018).

To address the above challenges in orthology prediction, we developed KinOrtho as a complementary approach for efficient and accurate identification of human kinase orthologs across ~17,000 species, extending well beyond the 15 model organisms defined in the seminal study of Manning and colleagues (Manning et al. 2002) and a recently updated kinase-centric database with kinases from 2,000 species (Krupa et al. 2004). KinOrtho is query-based and achieves increased sensitivity by combining similarities in the commonly conserved protein kinase domain and flanking regulatory domains. This enables us to develop one-to-one orthology relationships providing a finer resolution of orthologs across species compared to previous efforts. We apply KinOrtho towards identifying putative proteins involving fusion or fission events, or so-called “Rosetta Stone protein” (Marcotte et al. 1999). By integrating evolutionary information with gene expression patterns, we identify a potential functional association between an understudied kinase “Serine/threonine-protein kinase TAO1”

(TAOK1) and a metabolic kinase “Phosphatidylinositol-4-phosphate 3-kinase” (PI3KC2A) in autophagy. Using KinOrtho-defined orthologs, Gene Ontology (GO) annotations, and machine learning models, we prioritize understudied kinases for functional studies by developing a Novel Inferred Annotation Score (NIAS). The KinOrtho pipeline and ortholog datasets are available at the GitHub (https://github.com/esbgkannan/KinOrtho), and the patterns of conservation in aligned orthologs sequences are visualized in both KinView (McSkimming et al. 2016) (https://prokino.uga.edu/kinview/) and Pharos (Sheils et al. 2021) (https://pharos.nih.gov/).

## Results

### Overview of KinOrtho algorithm

KinOrtho is a query- and graph-based orthology inference method that combines full-length and domain-based orthology inference approaches. It consists of two pipelines (full-length and domain-based) and six main steps: (i) homology search, (ii) building kinome databases, (iii) all-vs-all homology search, (iv) orthology inference, (v) cluster analysis, and (vi) combining the results from two pipelines (Figure 1).

**Figure 1.**
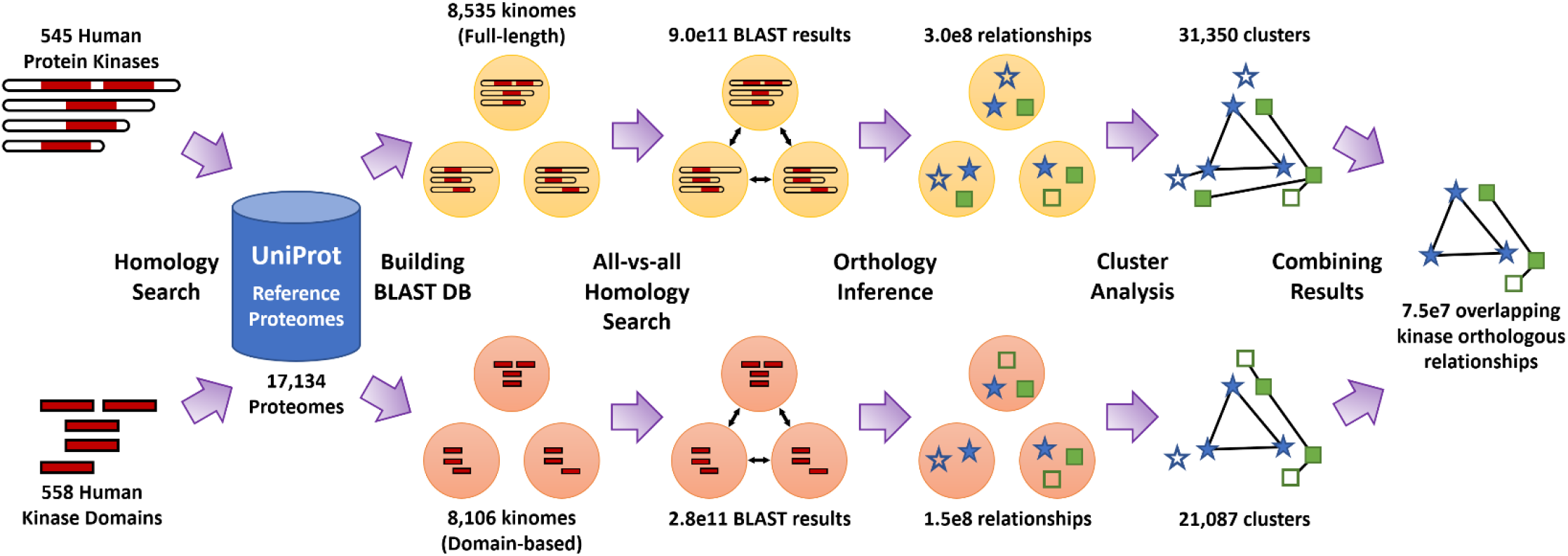
Overview of the KinOrtho algorithm. KinOrtho is an orthology inference method combining full-length and domain-based approaches. It consists of six main steps: (i) homology search against reference proteomes, (ii) building BLAST databases, (iii) all-vs-all homology search, (iv) orthology inference, (v) cluster analysis, and (vi) combining the results of full-length and domain-based approaches.

Because KinOrtho is query-based, it omits a large portion of unnecessary sequence comparisons unrelated to the query sequence(s). This characteristic makes KinOrtho a more efficient tool to identify orthologs of interest across the tree of life. We applied KinOrtho to identify the orthologs of 545 human kinases across some 17,000 species found in UniProt reference proteomes (Suzek et al. 2007). Without target genes, traditional orthology inference methods start from an all-vs-all homology search, which would require orders of magnitude (more than two quadrillion) pairwise sequence comparisons for this sample of reference proteomes. Because our query sequences were human kinases, only about eight thousand species were found to have human kinase homologs, which resulted in a nearly 2,000-fold reduction in the number of comparisons to be made (Supplementary Table S1). This makes KinOrtho one of the most computationally efficient orthology inference methods currently available for the identification of kinases.

When performing orthology inference, we adopted the definition of orthologous relationship used by OrthoMCL (Li et al. 2003) (see Methods), which resulted in twice as many orthologous relationships using similarity in full-length sequences compared to similarities within the kinase domain alone (302 million for full-length vs. 148 million for kinase domain; Supplementary Table S1). However, the application of graph-based clustering and further refinement of the clusters resulted in a comparable number of orthologous relationships in the full-length (97 million) and domain-based (100 million) pipelines (referred to as full-length set and domain-based set, respectively). Finally, the combination of both pipelines resulted in 75 million overlapping orthologous relationships (termed “overlapping set” from here on), including ~133,000 relationships between human kinases and kinases from other species (Supplementary Table S1). Since this is the most refined set of relationships obtained from KinOrtho, this overlapping set will be referred to as KinOrtho throughout this manuscript. In contrast, the reference to the full-length and domain-based results will be stated explicitly when they occur.

### Benchmarking and comparison of KinOrtho with other orthology inference methods

To evaluate and compare the performance of KinOrtho with the ability of other orthology inference methods in identifying kinase orthologs, we applied KinOrtho to the well-curated Quest for Orthologs (QfO) reference proteomes 2018 (Altenhoff et al. 2016). As shown in Figure 2a, the overall comparison metrics for KinOrtho are better (in terms of the overall precision and recall) than existing methods in the benchmarking datasets based on the enzyme classification conservation test, agreement with reference gene phylogenies, and species tree discordance benchmarks. The remaining metrics are shown in Supplementary Figure S1. It is also important to note that the selection of orthologs from KinOrtho’s full-length pipeline, domain-based pipeline, and the overlapping results all yielded similar performance (compared to other methods), suggesting robustness and agreement across these methods.

**Figure 2.**
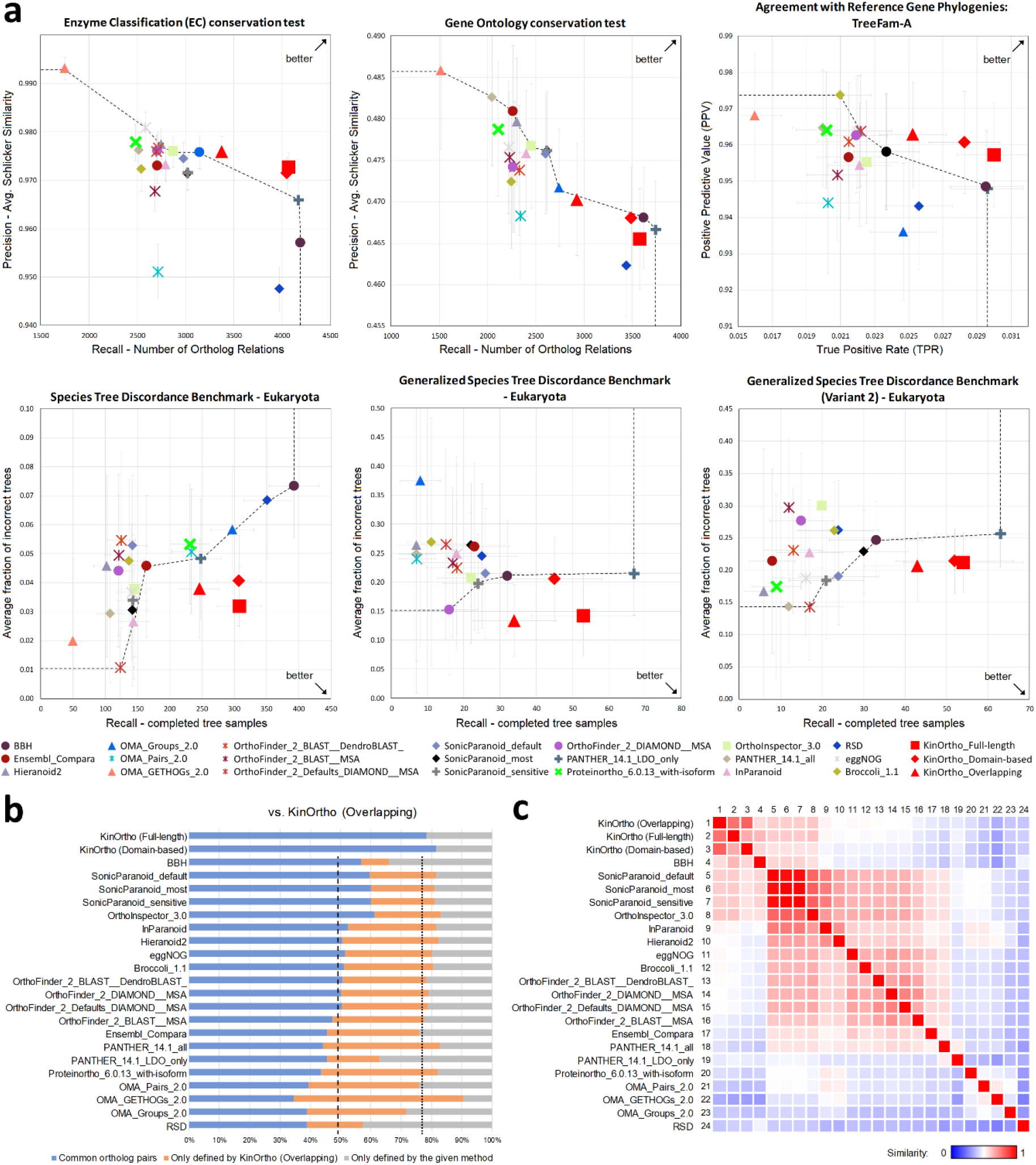
Benchmarking and comparison of KinOrtho with other methods. (a) Evaluations of the kinase orthologs identified by KinOrtho (marked in red) and 21 other methods. The title of each plot represents the evaluation metric. The dotted line represents the Pareto frontier, which runs over the participants with the best efficiency (except KinOrtho). The arrow in the plot shows the optimal corner. Red square: KinOrtho full-length set; red diamond: KinOrtho domain-based set; red triangle: KinOrtho overlapping set. (b) The 100% stacked bar chart shows the overlap in kinase orthologs identified by KinOrtho versus other orthology inference methods (blue region); a dashed line indicates the average percentage of the overlaps (KinOrtho Full-length and KinOrtho Domain-based do not count). The orange region represents the percentage of orthologs only identified by KinOrtho; a dotted line indicates the average percentage of the blue and orange regions. The gray region shows the percentage of orthologs unique to the compared method. (c) The heat map represents the Jaccard similarity matrix among orthology inference methods. Method indices are shown on the left and top of the matrix.

Next, a direct comparison between the pairs of orthologs identified by KinOrtho was performed alongside other methods in order to gain insights into overlapping predictions. In general, 35-60% of the orthologs identified by KinOrtho were also identified by other methods (blue bars in Figure 2b). Besides, KinOrtho (overlapping set) consistently found orthologs not identified by other methods (orange bars in Figure 2b). Several unique KinOrtho-defined human kinase orthologs, such as cyclin-dependent protein kinase (CDK) orthologs, are described in Supplementary Results and shown in Supplementary Figure S2-S4. On the other hand, KinOrtho consistently omitted at least 10% of the orthologs (average: 23.2%; gray bars in Figure 2b) that were identified by other methods. This number is significantly reduced when considering KinOrtho full-length or domain-based sets alone (average: 14.3% and 17.2%, respectively; gray bars in Supplementary Figure S5), suggesting that KinOrtho eliminates putative ortholog sequences that lack the well-defined bilobal kinase domain associated with protein kinases. The ability of KinOrtho to delineate the orthologs based on the protein kinase domain against the orthologs based on other conserved domains is described in Supplementary Results and shown in Supplementary Figure S6. Additional details about the utility and benchmarking of the domain-based approach are discussed below. Finally, we generated a similarity heat map to quantify orthology predictions by KinOrtho and other methods (Figure 2c). Similarities measured by the Jaccard similarity coefficient between two ortholog sets ranged from 25.3% (Reciprocal Smallest Distance (RSD) (Wall et al. 2003) vs. Orthologous Matrix (OMA) (Train et al. 2017) to 81.5% (SonicParanoid (Cosentino and Iwasaki 2019) vs. OrthoInspector (Nevers et al. 2019). The average similarity among all methods was 50.4%. Orthology results from Bidirectional Best Hits (BBH) (Overbeek et al. 1999) and two BBH- and graph-based methods, OrthoInspector and SonicParanoid, were found to have the most agreement with KinOrtho results (average similarities: 59.9%, 59.8%, and 58.7%, respectively).

### Inferring functional associations using KinOrtho-based identification of kinase domain fusion and fission events

KinOrtho’s ability to find orthologs for individual domains allows identifying domain fusion and fission events for kinases with multiple kinase domains. In the human kinome, there are 13 kinases with two tandem kinase domains within the same polypeptide, many of which are functionally annotated phosphorylation targets of Mitogen Activated Protein Kinase (MAPK) signaling pathways. Figure 3a illustrates the four scenarios of finding domain-based orthologs for these 13 kinases: (1) tandem domains in one kinase match tandem domains in another kinase, (2) tandem domains in one kinase match tandem domains in another kinase in reverse order, (3) two domains from two human kinases match tandem domains in one kinase from another species, and (4) tandem domains in one human kinase match two domains from two kinases in another species. Traditional full-length BBH-based orthology inference methods do not have the resolution to distinguish between these scenarios. However, KinOrtho’s domain-based approach allows the definition of orthologs from all scenarios, thus identifying fusion and fission events in orthologous sequences.

**Figure 3.**
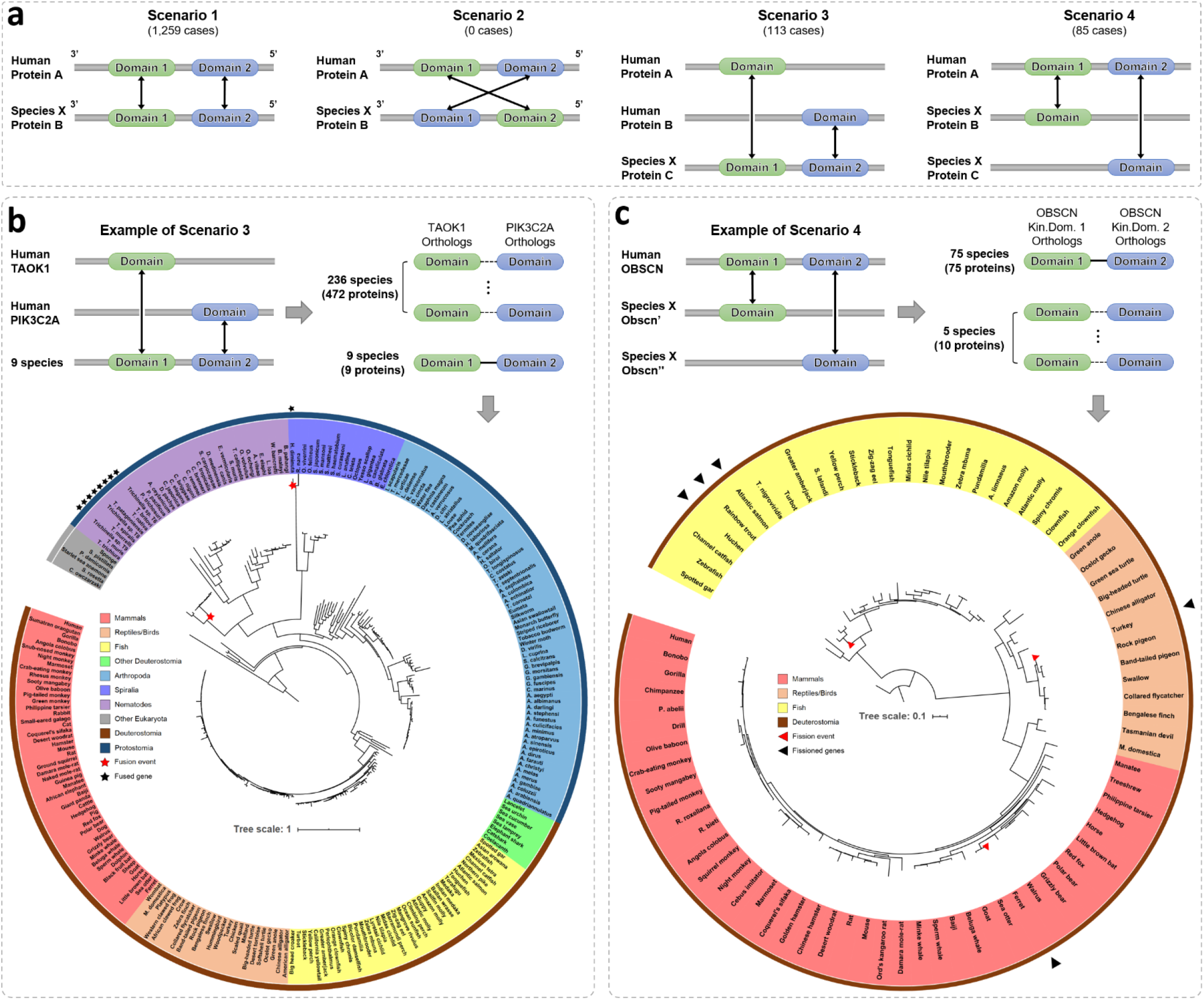
Scenarios of a single protein with two kinase domains and examples of potential gene fusion and fission events. (a) Four scenarios of a single protein with two kinase domains and their domain-based orthologs. Arrows represent orthologous pairs. The number of cases for each scenario is shown in parentheses. (b) Example of Scenario 3 and phylogenetic tree analysis on the fusion event of TAOK1 orthologs and PIK3C2A orthologs. Human TAOK1’s and PIK3C2A’s domain-based orthologs in 245 species were aligned, concatenated (represented by a dotted line if these two domains are from different species), and used to build a phylogenetic tree. Species names are labeled at the leaves of the circular-mode phylogenetic tree. The leaves are colored according to the clade of the species (refer to legend). Black stars mark the species with a TAOK1-PIK3C2A fused gene. The time when potential fusion events occurred is indicated by the red stars shown on the tree. (c) Example of Scenario 4 and analysis of the fission event of OBSCN orthologs. The domain-based orthologs of human OBSCN’s two kinase domains in 80 species were used to build a phylogenetic tree. Black triangles mark the species with fissioned genes. Red triangles indicate the time when potential fission events occurred.

Scenario 3 in Figure 3a reflects potential domain fusion events. Although we identified 113 potential fusion events, we use TAOK1 and PIK3C2A as an example to illustrate how integrating evolutionary data with other contextual data (protein-protein interaction, co-expressions, and co-occurrence) can reveal potential functions for understudied kinases (Supplementary Data S1). TAOK1, an understudied kinase, belongs to the STE20 family, while PIK3C2A belongs to the PI3K family. We found nine kinases with two domains matching both the human TAOK1 kinase domain and PIK3C2A kinase domain (Figure 3b). In addition, we identified 236 species with both TAOK1 and PIK3C2A domain-based orthologs in different kinases. We concatenated the sequences of TAOK1 orthologs and PIK3C2A orthologs for each species and then built a phylogenetic tree. We found that eight kinases with TAOK1 and PIK3C2A domains reside in the same clade of Nematodes. Based on this, we postulate two potential fusion events (indicated by red stars on the phylogenetic tree, Figure 3b). Proteins involved in a fusion event usually belong to the same functional category (Yanai et al. 2001). As an example, TAOK1, an understudied kinase, shows high co-expression with PIK3C2A in 17,382 normal samples and 1,376 cancer samples in the Genotype-Tissue Expression project (Consortium 2013) (GTEx, version 8) and the Cancer Dependency Portal (Tsherniak et al. 2017) (DepMap, 20Q4), respectively. The correlations (Pearson correlation coefficient = 0.856 in normal samples and 0.612 in cancer samples) are among the top 0.15% of all kinase pairs (Supplementary Figure S7). The co-expressed patterns are conserved in *A.aegypti, B.taurus, D.melanogaster*, and *S.mansoni* (STRING (Szklarczyk et al. 2019), version 11.0), suggesting a possible physical interaction. Moreover, TAOK1 and PIK3C2A have been reported to be involved in the autophagy response (Bennetzen et al. 2012). Based on the finding of these evolutionary fusion events and contextual data, we predict a functional association between the understudied kinase TAOK1 and PIK3C2A in human cellular biology, perhaps involving communication between the membrane, where phospholipids are sensed, and the cytosol, where TAOK1 has known functions in relaying information to MAPK pathways.

We also analyzed cases in Scenario 4 for potential domain fission events. We found ten kinases in five species (ferrets, turkeys, Atlantic salmon, rainbow trout, and huchen) matching the tandem kinase domains in human Obscurin (OBSCN) kinase (Figure 3c). Besides, the tandem kinase domain arrangement in OBSCN is conserved in 75 species (Scenario 1). In species where the tandem domains are encoded in two different proteins, we concatenated the domains and performed phylogenetic comparisons with species where the two domains are naturally fused. The concatenated sequences of Atlantic salmon, rainbow trout, and huchen occur in the same clade. Based on the phylogenetic tree, we estimate three kinase domain fission events (marked by red triangles in Figure 3c). Although the functional significance of these fission events is unclear, the established role of OBSCN in eye development (Kontrogianni-Konstantopoulos and Bloch 2005; Perry et al. 2013) suggests a role for these events in the evolution of vision in these species (Dickson 1992; Fraser et al. 1993; Kadri et al. 1997; Fox and Marini 2014).

### Phylogenetic profile analysis reveals the evolutionary depth of human protein kinase conservation and enriched molecular functions across species

We next sought to classify human kinases based on conservation depth across species by building a phylogenetic profile of KinOrtho-defined orthologs. Figure 4a highlights a human kinase phylogenetic profile consisting of 558 human kinase domains and their orthologs across 561 clades. As expected, human kinase orthologs are barely present in bacteria, archaea, and viruses, except for the orthologs of eukaryotic-like protein kinases. Consistent with previous findings, four eukaryotic-like kinases, Protein adenylyltransferase SelO, mitochondrial (SELENOO; 3,936 orthologs), AarF domain-containing protein kinase 1 (ADCK1; 3,258 orthologs), Ketosamine-3-kinase (FN3KRP; 2,234 orthologs), and Serine/threonine-protein kinase RIO1 (RIOK1; 1,849 orthologs) have the most orthologs. In contrast, Casein kinase II subunit alpha 3 (CSNK2A3), Rhodopsin kinase GRK1 (GRK1), Putative serine/threonine-protein kinase PRKY (PRKY), and Probable serine/threonine-protein kinase SIK1B (SIK1B) are “orphan” kinases with no orthologs based on KinOrtho’s stringent criteria. The phylogenetic profile also shows that the kinases in tyrosine kinase (TK), tyrosine kinase-like (TKL), and receptor guanylate cyclase (RGC) groups are mainly conserved in Metazoa (including mammals, reptiles, birds, fish, and protostomes), which is consistent with the findings of a previous study (Suga et al. 2012).

**Figure 4.**
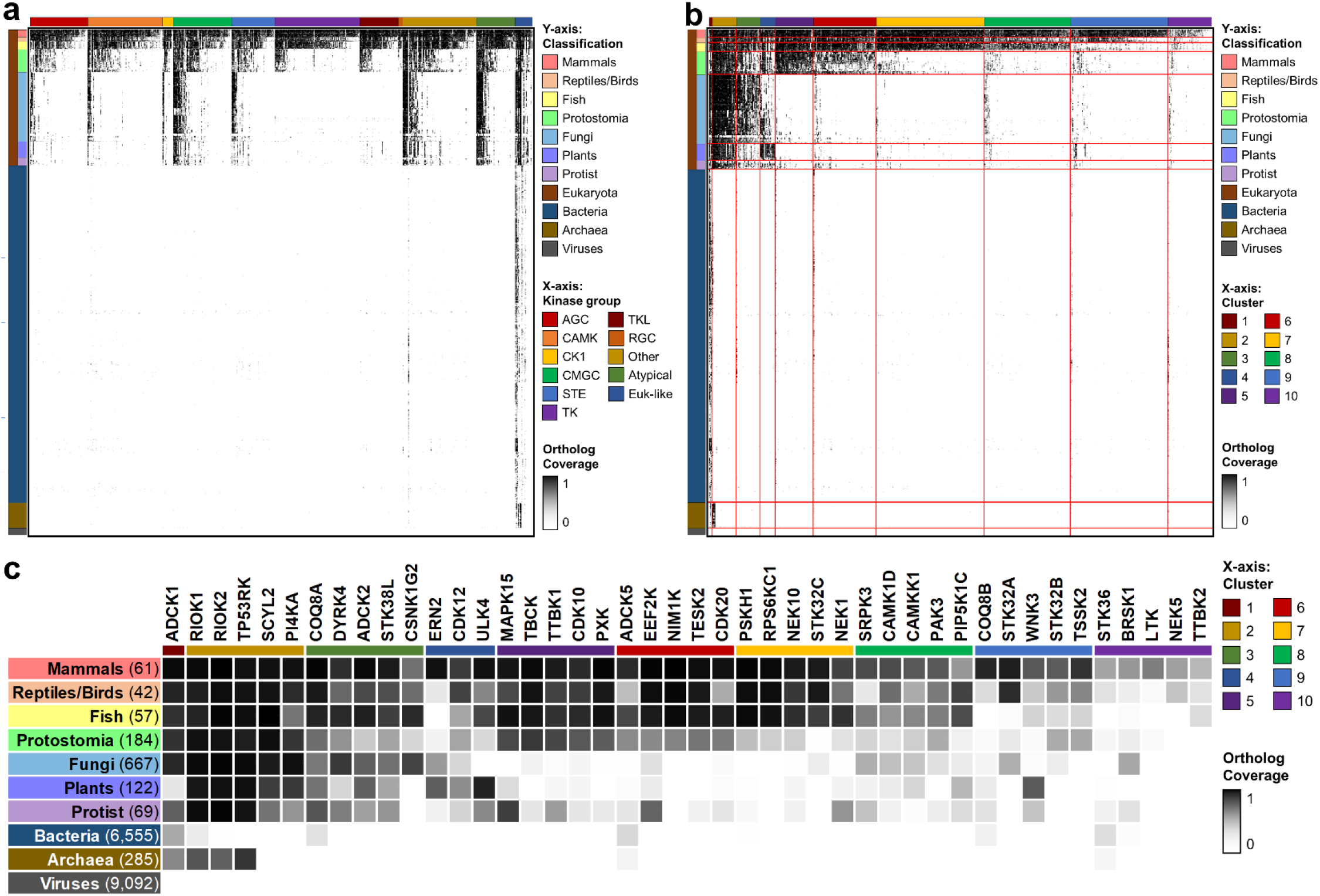
Cluster analysis of the phylogenetic profile of human kinases. (a) Phylogenetic profile of human kinases. X-axis: 558 human kinase domains, ordered by the kinase group and the number of orthologs identified by KinOrtho. Y-axis: 561 clades, ordered by classification. Each dot’s color represents the coverage of the human kinase ortholog in each clade. (b) Phylogenetic clusters of human kinases. The human kinase domains in the x-axis are in the order of clusters and the number of orthologs. The boundaries of each cluster and classification are highlighted in red. (c) Top 5 understudied kinases with the most orthologs in each cluster. The number of species in each classification is shown in parentheses.

Many understudied kinases have escaped analysis due to weak conservation in model organisms. Based on the distribution of orthologs across different species, we organized human kinases into 10 clusters (Figure 4b). The top 5 understudied kinases with the most orthologs in each cluster are highlighted in Figure 4c. Kinases within each cluster are closely related (small Euclidean distance) with high co-occurrence with each other. Using this clustered phylogenetic profile, we sought to identify potentially conserved kinase-regulated biological functions across species. For example, because the kinases in Cluster 5 are highly conserved in Metazoa, we can hypothesize a role for these kinases in metazoan-specific biological functions. To this end, we performed Gene Ontology (GO) enrichment analyses using the GO annotations of all kinases as background. We identified 802 significantly enriched GO terms (false discovery rate (FDR) < 0.05), and the GO term is annotated for at least five human kinases in the cluster. The top three enriched GO terms for each cluster are shown in Table 1, and the entire list is shown in Supplementary Data S2.

**Table 1.** GO term enrichment analysis on the phylogenetic clusters of human kinases. Based on fold enrichment, only the top three enriched GO terms in each cluster are shown. Cluster: the cluster ID shown in Figure 4b; #Kinases: the total number of human kinases and their orthologs in the cluster; K: the total number of kinases associated with the GO term; k: the number of kinases associated with the GO term in the cluster; Fold: fold enrichment; FDR: false discovery rate.

| Cluster | #Kinases | Conserved in | Enriched GO term | K | k | Fold | FDR |
| --- | --- | --- | --- | --- | --- | --- | --- |
| 2 | 27,128 | Eukaryota | GO:0000075<br>(Cell cycle checkpoint) | 4,765 | 3,619 | 3.66 | < 1.0E-320 |
|  |  |  | GO:0098805<br>(Whole membrane) | 1,289 | 890 | 3.33 | 2.3E-306 |
|  |  |  | GO:0046488<br>(Phosphatidylinositol metabolic process) | 2,634 | 1,772 | 3.24 | < 1.0E-320 |
| 3 | 16,899 | Vertebrata,<br>Protostomia<br>(partial),<br>Fungi | GO:0000165<br>(MAPK cascade) | 1,943 | 1,025 | 4.08 | < 1.0E-320 |
|  |  |  | GO:0005694<br>(Chromosome) | 284 | 120 | 3.27 | < 1.0E-320 |
|  |  |  | GO:1902749<br>(Regulation of cell cycle G2/M phase transition) | 983 | 380 | 2.99 | 2.2E-10 |
| 4 | 6,933 | Vertebrata,<br>Protostomia<br>(partial),<br>Plants | GO:0010468<br>(Regulation of gene expression) | 9,690 | 796 | 1.55 | < 1.0E-320 |
|  |  |  | GO:0006807<br>(Nitrogen compound metabolic process) | 20,390 | 1,470 | 1.36 | < 1.0E-320 |
|  |  |  | GO:0044238<br>(Primary metabolic process) | 23,630 | 1,663 | 1.33 | 2.1E-09 |
| 5 | 12,496 | Metazoa | GO:0008047<br>(Enzyme activator activity) | 778 | 341 | 4.59 | 1.9E-10 |
|  |  |  | GO:0090287<br>(Regulation of cellular response to growth factor stimulus) | 627 | 249 | 4.15 | 3.7E-11 |
|  |  |  | GO:0016055<br>(Wnt signaling pathway) | 415 | 161 | 4.06 | < 1.0E-320 |
| 6 | 16,115 | Vertebrata,<br>Protostomia<br>(partial) | GO:0045669<br>(Positive regulation of osteoblast differentiation) | 437 | 243 | 4.51 | 1.1E-102 |
|  |  |  | GO:0030500<br>(Regulation of bone mineralization) | 508 | 244 | 3.90 | < 1.0E-320 |
|  |  |  | GO:0034645<br>(Cellular macromolecule biosynthetic process) | 797 | 330 | 3.36 | 1.0E-10 |
| 7 | 17,842 | Vertebrata | GO:0042629<br>(Mast cell granule) | 54 | 54 | 7.33 | 5.5E-46 |
|  |  |  | GO:0030522<br>(Intracellular receptor signaling pathway) | 152 | 146 | 7.04 | 8.6E-116 |
|  |  |  | GO:0042102<br>(Positive regulation of T cell proliferation) | 211 | 202 | 7.01 | 1.6E-159 |
| 8 | 10,994 | Mammals,<br>Reptiles/Birds<br>(partial),<br>Fish | GO:0051965<br>(Positive regulation of synapse assembly) | 250 | 177 | 8.42 | 2.2E-128 |
|  |  |  | GO:0005004<br>(GPI-linked ephrin receptor activity) | 215 | 130 | 7.19 | 5.4E-81 |
|  |  |  | GO:0005005<br>(Transmembrane-ephrin receptor activity) | 267 | 143 | 6.37 | 3.9E-79 |
| 9 | 10,202 | Mammals,<br>Reptiles/Birds<br>(partial) | GO:0001669<br>(Acrosomal vesicle) | 103 | 75 | 9.33 | 3.2E-58 |
|  |  |  | GO:0031253<br>(Cell projection membrane) | 283 | 161 | 7.29 | 1.8E-99 |
|  |  |  | GO:0014068<br>(Positive regulation of phosphatidylinositol 3-kinase signaling) | 413 | 164 | 5.09 | 4.8E-11 |
| 10 | 3,043 | Mammals<br>(partial) | GO:0035173<br>(Histone kinase activity) | 665 | 96 | 6.20 | 1.1E-11 |
|  |  |  | GO:0050321<br>(Tau-protein kinase activity) | 318 | 43 | 5.81 | 8.7E-12 |
|  |  |  | GO:0051051<br>(Negative regulation of transport) | 818 | 98 | 5.15 | < 1.0E-320 |

The orthologs of human kinases in Cluster 2 are present in most eukaryotic species. The most enriched GO term in Cluster 2 is a biological process term “cell cycle checkpoint” (GO:0000075), which encompasses a variety of DNA and spindle-assembly checkpoints, well-established control mechanisms that control progression through the eukaryotic cell cycle (Hartwell and Weinert 1989). EKC/KEOPS complex subunit TP53RK (TP53RK), an understudied kinase with 1,367 orthologs, plays a vital role in the cell cycle and G1 checkpoint control (Abe et al. 2001; Richardson et al. 2012). However, this GO term is currently absent in both human TP53RK annotation and TP53RK ortholog annotation. Kinases in Cluster 9 are mostly present in mammals. Consistently, the cellular component term “acrosomal vesicle” (GO:0001669) is the most enriched GO term in Cluster 9. Acrosomal vesicles, components in the sperm’s head, contain enzymes essential for fertilization (Moreno et al. 2000). All members of testis-specific serine/threonine-kinases (TSSK) belong to Cluster 9, and they are all understudied kinases: TSSK1B, TSSK2, TSSK3, TSSK4, and TSSK6. Currently, TSSK1B, TSSK2, and TSSK4 are annotated with this GO term. Although TSSK3, TSSK6, and their orthologs lack this annotation, both TSSK3 and TSSK6 are highly expressed in testis (median Transcripts Per Million = 60.86 and 424.6, respectively) (Consortium 2013). TSSK6 is also reported to be involved in the acrosome reaction and egg fertilization (Sosnik et al. 2009). Therefore, based on the cluster analysis of the phylogenetic profile, we predict TSSK3 and TSSK6 function in acrosomal biology and vesicle localization.

### Machine learning model to prioritize understudied kinases using KinOrtho and GO annotations

The human kinome contains several understudied kinases of unknown function. We next wanted to investigate if KinOrtho-defined orthologs, along with sequence similarities and GO annotations from different species, can be used to infer the functions of understudied kinases using “guilt-by-association” (Oliver 2000) and machine learning methods. To this end, we trained machine learning classifiers using orthology and functional annotations of well-studied human kinases to predict whether functional annotation could be transferred from orthologs in other species to human kinases. In brief, we built training sets using the manually curated GO annotations of well-studied kinases (Figure 5). The input features of each training instance represent the GO annotation status of human kinase orthologs; the output shows whether the human kinase has this GO term annotation or not. Input features were weighted based on the sequence similarities between the human kinase and orthologs. The training sets contained 0.3 million instances with 730 GO terms for 393 well-studied kinases and their orthologs across 176 species. After class balancing and 10-fold cross-validation, random forest displayed high prediction accuracies among various machine learning methods attempted (90.9%, 92.1%, and 95.5% for the training sets of biological process, cellular component, and molecular function, respectively; Table 2).

**Figure 5.**
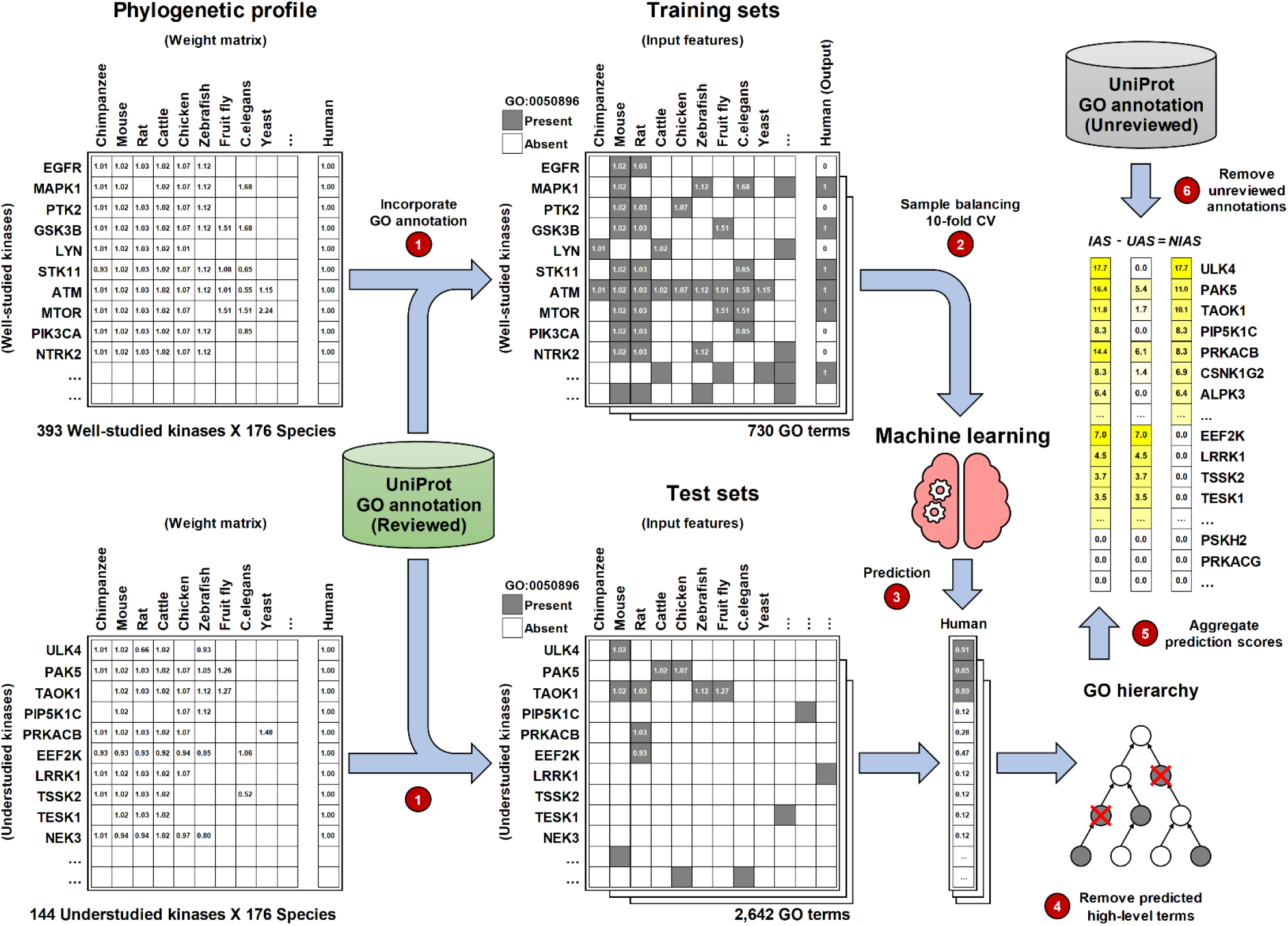
Calculating the Novel Inferred Annotation Score (NIAS) for understudied kinases using the phylogenetic profile, GO annotations, and machine learning methods. The weight matrices represent the normalized sequence similarities between the human kinase and orthologs. The training sets and test sets show an example of a GO term (GO:0050896, response to stimulus) annotation status across well-studied human kinases, understudied human kinases, and orthologs. CV: cross-validation; IAS: Inferred Annotation Score; UAS: Unreviewed Annotation Score; NIAS: Novel Inferred Annotation Score.

**Table 2.** Performance of GO annotation prediction in each training set. The best performance in each measurement and each GO domain is highlighted in bold.

| GO domain | Model | Accuracy | Precision | Recall | F-measure | AUC |
| --- | --- | --- | --- | --- | --- | --- |
| Biological process | Logistic regression | 0.896 | 0.942 | 0.844 | 0.891 | 0.914 |
|  | SVM | 0.896 | 0.943 | 0.843 | 0.890 | 0.896 |
|  | Random forest | **0.909** | **0.944** | **0.869** | **0.905** | **0.923** |
| Cellular component | Logistic regression | 0.908 | 0.955 | 0.856 | 0.903 | 0.924 |
|  | SVM | 0.908 | 0.955 | 0.858 | 0.903 | 0.908 |
|  | Random forest | **0.921** | **0.959** | **0.880** | **0.918** | **0.932** |
| Molecular function | Logistic regression | 0.940 | 0.959 | 0.919 | 0.939 | 0.961 |
|  | SVM | 0.942 | 0.959 | 0.925 | 0.941 | 0.942 |
|  | Random forest | **0.955** | **0.964** | **0.945** | **0.955** | **0.965** |

Next, we used the trained models (random forests) to predict the functions of understudied kinases. To this end, we constructed test sets in which understudied human kinases could be annotated based on the GO terms available for one or more of their orthologs. This resulted in 16 thousand instances with 2,642 GO terms for 144 understudied kinases. Application of the pre-trained random forest models on the test sets resulted in 11,573 predictions of kinase-GO term pairs as present (Supplementary Table S2). Among these predicted annotations, 8,933 predictions (77.2%) already existed in the UniProt as manually reviewed annotations, while 2,640 predictions (22.8%) were not found in the UniProt. Instead of referring to these 2,640 predictions as false positives, we considered them as missing annotations. In fact, 1,452 of them (55%) were found to be unreviewed electronic annotations from Ensembl (Yates et al. 2020), InterPro (Mitchell et al. 2019), the UniProt Consortium, or the GO Consortium. The remaining 1,188 annotations, including 236 lowest-level GO term annotations, were novel inferred annotations (available in Supplementary Data S3). By aggregating the prediction score of each novel inferred annotation, we calculated a Novel Inferred Annotation Score (NIAS) for each understudied human kinase (the last step in Figure 5).

Our analysis reveals that Serine/threonine-protein kinase ULK4 (ULK4) has the highest NIAS among all understudied human kinases. It has 22 novel inferred annotations. Fifteen of them with prediction scores higher than 0.9 are inferred from mouse/rat Ulk4. Twelve of these inferred annotations with a high score are associated with neuronal function and brain development, such as “ventricular system development” (GO:0021591), “corpus callosum development” (GO:0022038), “neuronal stem cell division” (GO:0036445), and “GABAergic neuron differentiation” (GO:0097154). A role for human ULK4 in neuronal function and brain development has been suggested (Lang et al. 2014; Lang et al. 2016; Khamrui et al. 2020), and it is an unusual pseudokinase that binds to nucleotides in the absence of cations (Murphy et al. 2014; Eyers 2020). Serine/threonine-protein kinase PAK 5 (PAK5), an understudied kinase with the second highest NIAS, has 12 novel inferred annotations. The association between PAK5 and “activation of MAPK activity” (GO:0000187), the GO term with the highest prediction score for PAK5, is also known from the literature, where these kinases act as upstream regulators of MAPK modules (Pandey et al. 2002). We also identified 80 understudied kinases with a NIAS of 0. We can still prioritize these proteins by Inferred Annotation Score (IAS, which aggregates all prediction scores regardless of existing unreviewed annotations; see Methods) for further manual curation or experimental validation. For example, the NIAS of Eukaryotic elongation factor 2 kinase (EEF2K) is 0. Kinases in this list include pseudokinases such as Serine/threonine-protein kinase H2 (PSKH2), which represent the “darkest” of kinases with little or no information across species and no functional biology currently reported (Shrestha et al. 2020a).

## Discussion

Here we map human kinase orthologs across diverse species by developing a kinase orthology inference method called KinOrtho. We demonstrate that KinOrtho performs better than existing orthology inference methods based on comparisons across standard benchmarking datasets and metrics. KinOrtho utilizes domain-based orthology inference to eliminate orthologs with no kinase domains, allowing researchers to focus on the functional domains of interest. KinOrtho’s query-based characteristic enables users to identify orthologs of specific kinases across thousands of species within a reasonable time. In contrast to orthologous groups provided by other methods, this approach provides one-to-one ortholog, in-paralog, and co-ortholog relationships, thereby revealing functional relationships and separating even the most closely related paralogous sequences.

While KinOrtho’s performance is better than existing methods based on metrics in the benchmarking dataset (Figure 2a and Supplementary Figure S1), overlap in orthologous relationships (not only kinase orthologs) defined by various methods in the benchmarking dataset is significantly low (only 29.9% similarity; Supplementary Figure S8), presumably because of the variability in orthology definition, methods used, or even potential genome assembly errors in the UniProt reference database. Thus, the interpretation of ortholog sets should be made with some caution. The Alliance of Genome Resources (AGR) has recently established orthologous relationships among humans and six model organisms: *Caenorhabditis elegans, Drosophila melanogaster*, *Danio rerio*, *Mus musculus*, *Rattus norvegicus*, and *Saccharomyces cerevisiae* (Alliance of Genome Resources 2020). The orthologous relationships in AGR are based on the consensus of seven orthology inference methods (Thomas et al. 2003; Kaduk et al. 2017; Train et al. 2017; Emms and Kelly 2019; Nevers et al. 2019; Yates et al. 2020) and five databases (Povey et al. 2001; Li et al. 2006; DeLuca et al. 2012; Huerta-Cepas et al. 2014; Ruzicka et al. 2019). Comparison of KinOrtho-defined human kinase orthologs with AGR-defined orthologs reveals nearly 70.5% similarity. The greatest difference in kinase orthology sets occurs in the CMGC and CAMK groups, presumably because of the deeper conservation of these kinases across taxa (Supplementary Figure S2).

In our previous study (Huang et al. 2018), we developed an annotation score (AS) for prioritizing understudied kinases based on existing knowledge; in this study, we propose a complementary NIAS for prioritizing understudied kinases based on missing knowledge. Together, these scores provide a comprehensive metric for pursuing understudied kinase studies. To accurately infer the function of understudied kinases using orthologous relationships, extensive annotations from broad coverage of species are needed. For illuminating understudied kinases, or understudied enzyme superfamilies, ion channels, or G-protein-coupled receptors, a broader collection of manually curated biological functions from various species would be immensely helpful.

In conclusion, we have developed an efficient query-based orthology inference method that combines full-length and domain-based orthology inference methods to comprehensively map human kinase orthologs across the tree of life in this study. KinOrtho performed better than existing methods in a benchmarking dataset and identified putative domain fusion and fission events. We confirmed kinase-associated molecular functions enriched across species using phylogenetic profiles after identifying overlapping orthologous relationships from full-length and domain-based pipelines. Finally, we prioritized and inferred functions of understudied human kinases using KinOrtho-defined orthology and GO annotations as features in machine learning. Our studies serve as a conceptual starting point for investigating understudied human kinase biology by leveraging evolutionary information. This is exemplified, but by no means limited-to, pharmacologically tractable enzyme superfamilies such as the protein kinases.

## Methods

### KinOrtho workflow

KinOrtho is a query-based, graph-based, and combinatorial orthology inference method. It consists of six main steps (Figure 1):

1. Homology search for the query sequences of interest against reference proteomes
2. Building Basic Local Alignment Search Tool (Altschul et al. 1990) (BLAST) databases, containing full-length and domain-based databases
3. All-vs-all homology search for the rebuilt databases
4. Orthology inference and determining orthologs, paralogs, and co-orthologs
5. Cluster analysis and filtering out the orthologous relationships between two proteins in different clusters or the clusters without query sequences
6. Combining the results of full-length and domain-based methods

#### Query sequences

This study’s query sequences are based on a recent article that re-analyzes the human kinome’s composition (Moret et al. 2020). We collected 545 human kinases, containing 483 protein kinases, 19 eukaryotic-like protein kinases, and 43 atypical protein kinases. Based on a manually curated eukaryotic protein kinase sequence profile (Kwon et al. 2019), Pfam (El-Gebali et al. 2019), and Conserved Domain Database (Lu et al. 2020), we manually annotated and collected 558 kinase domain sequences from the 545 human kinases. More information about the domain name, domain boundary, and kinase group are available in Supplementary Data S4.

#### Reference proteomes

We applied KinOrtho to the UniProt reference proteomes (release 2019_11), which are chosen to broadly represent the taxonomic diversity (Suzek et al. 2007). It is also the most well-curated and extensive collection of entire proteomes across the tree of life. The reference proteomes contain 18,870,318 protein sequences spanning the tree of life (Supplementary Table S1). To benchmark the performance of KinOrtho, we applied KinOrtho to the Quest for Orthologs (QfO) reference proteomes 2018 (Altenhoff et al. 2016), which contains 885,338 protein sequences from 48 eukaryotic species, 82,507 sequences from 23 bacterial species, and 17,317 sequences from 7 archaea species (Supplementary Table S1). To benchmark KinOrtho based on a domain-based kinase classification, we also applied KinOrtho to the model organisms in KinBase (Manning), which includes 15 species and 7,597 kinase sequences (Supplementary Table S3).

#### Homology search and building BLAST databases

Before performing a time-consuming all-vs-all homology search for all reference proteomes, KinOrtho looks for potential homologs of query sequences by screening the reference proteomes using NCBI BLAST+ (Camacho et al. 2009) (version 2.7.1) with default settings, except for the E-value threshold. Referring to other orthology inference methods, such as OrthoMCL-DB (Fischer et al. 2011) and PANTHER (Thomas et al. 2003), KinOrtho uses 10^−5^ as a default E-value threshold for BLAST search. This threshold has been demonstrated to balance between false-positive and false-negative rates (Chen et al. 2007). An additional experiment showed that choosing the default E-value threshold of BLAST+ (10^1^) yielded 19 times more sequence comparisons than choosing 10^−5^ in the benchmarking dataset (Supplementary Table S4) but reduced performance based on the six benchmarking metrics shown in Supplementary Figure S9. Then, KinOrtho builds two sets of BLAST databases as new reference proteomes (“kinomes” hereafter) based on full-length and domain-based query sequences. To build a full-length kinome for each proteome, KinOrtho keeps the sequences in the BLAST result, generates a new sequence file, and then applies the “makeblastdb” function provided by NCBI BLAST+. To build a domain-based kinome for each proteome, KinOrtho generates a new sequence file and builds a BLAST database based on the BLAST hit region (between “sstart” and “send”) of the sequences in the BLAST result. After building a set of full-length kinomes and a set of domain-based kinomes, KinOrtho performs an all-vs-all homology search for each set using the E-value threshold (10^−5^) mentioned above.

#### Orthology inference

The orthologous relationships identified by KinOrtho include orthologs, in-paralogs, and co-orthologs. KinOrtho defines a pair of one-to-one orthologs using the Bidirectional Best Hits (BBH) method (Overbeek et al. 1999). A pair of in-paralogs is defined as two protein sequences with a higher similarity score (BLAST bit score) in the same species than the homologous sequences in other species. A pair of co-orthologs is defined based on the following two criteria: (i) ortholog of one sequence is the in-paralog of the other, or (ii) in-paralog of each sequence are a pair of orthologs. Using all orthologous relationships as edges, KinOrtho builds two graphs by connecting the kinases in the full-length and domain-based kinomes, respectively.

#### Cluster analysis

To identify orthologous groups, KinOrtho performs the Markov Cluster (MCL) Algorithm (Van Dongen 2000) (version 14.137) for the two graphs. MCL is a fast, unsupervised clustering method using a simulation of flow in graphs. It has been utilized in other graph-based orthology inference methods (Li et al. 2003; Emms and Kelly 2015) and detecting protein families (Enright et al. 2002). In the orthologous relationship graphs, KinOrtho assigns the negative logarithm of the E-value as a weight for each edge. If an E-value is reported 0 by the BLAST program, KinOrtho assigns an arbitrary E-value of 10^−200^. Considering the systematic differences among species, such as nucleotide composition bias, KinOrtho normalizes the weights based on the method used by OrthoMCL (Li et al. 2003). For the orthologs or co-orthologs between any two species, KinOrtho normalizes the weights by dividing them by the average weight of all the orthologs or co-orthologs between the two species. For in-paralogs, the weights are divided by the average weight of all in-paralogs in each kinome. When performing MCL after setting a normalized weight for each edge, KinOrtho chooses 1.5 as a default inflation value to control the cluster tightness. This value is the best inflation value to balance the sensitivity and selectivity for functional classification (Li et al. 2003). Each protein is assigned to a cluster, after which KinOrtho refines orthologous relationships by filtering out the relationships between two proteins in different clusters or the clusters without query sequences.

#### Combining results

In the last step, KinOrtho combines the orthologous relationships from full-length and domain-based results. We define an “overlapping orthologous relationship” as a relationship present in both full-length and domain-based results. For example, in Scenario 1 of Figure 3a, if A_1_B_1_ (meaning the pair of A’s 1^st^ kinase domain and B’s 1^st^ kinase domain) and A2B2 are domain-based orthologs and A-B is a full-length ortholog pair, then both A_1_B_1_ and A_2_B_2_ are defined as overlapping orthologous relationships. However, in Scenario 4 of Figure 3a, if A_1_B_1_ and A_2_C_1_ are domain-based orthologs and A-B is a full-length ortholog pair, only A_1_B_1_ is an overlapping orthologous relationship, but A_2_C_1_ is not. Because non-overlapping relationships are also informative in domain-based orthology analyses, KinOrtho keeps all the results from full-length and domain-based methods.

### Comparison of orthology inference methods

There are 21 public orthology inference results available at Ortholog Benchmarking Webservice (Altenhoff et al. 2016) (Supplementary Table S5; similarity matrices are shown in Supplementary Figure S8). These datasets generated by full-length orthology inference methods contain the kinase relationships and all other proteins’ relationships in the QfO reference proteomes 2018. To make the orthologs identified by KinOrtho and those identified by the 21 methods comparable, we performed the following preprocessing for the compared datasets. First, because KinOrtho defines orthologs based on the BBH method, we only kept one-to-one relationships in the compared datasets. Second, to identify the kinase orthologs in the compared datasets, we only kept the relationships with at least one protein found in the ortholog relationships identified by KinOrtho (either full-length or domain-based approach). Finally, to identify human kinase orthologs in the compared datasets, we only kept the relationships involving human kinases. The numbers of remaining proteins and ortholog relationships are shown in Supplementary Table S5. We submitted these 21 preprocessed one-to-one kinase ortholog datasets to Ortholog Benchmarking Webservice for performance evaluation.

### Protein domain annotation

This study employed the annotations in Pfam (El-Gebali et al. 2019) (version 32.0) as known protein domain annotations. There are 305,472 proteins with at least one orthologous relationship identified by KinOrtho from the UniProt reference proteomes. In these proteins, 197,327 of them have 398,313 domain annotations, and 149,080 have at least one of the two major protein kinase domains: “Pkinase” and “Pkinase_Tyr”.

### Phylogenetic analysis

The phylogenetic tree analyses in this study were utilized to investigate the gene fusion and fission events of proteins with two kinase domains (Figure 3). First, we obtained the domain-based orthologs of the kinase domains of interest. To identify gene fusion events, we used TAOK1 and PIK3C2A as an example. There are 245 species (including humans) having both TAOK1 and PIK3C2A orthologs. To identify gene fission events, we used OBSCN as an example. There are 80 species with the orthologs of both OBSCN’s two kinase domains. Second, we aligned those two sets of kinase domain orthologs separately by Multiple Alignment using Fast Fourier Transform (Katoh et al. 2002) (MAFFT, version 7.407). We used options “L-INS-i”, “--localpair”, and maxiterate 10000” to generate more accurate alignments. Third, the two kinase domains’ alignments were concatenated as a single alignment file: the first kinase domain’s orthologs followed by the second kinase domain’s orthologs for each species. Fourth, we used IQ-TREE (Nguyen et al. 2015) with options “-m TEST” (standard model), “-bb 1000” (bootstrap replicates), and “-alrt 1000” (approximate likelihood ratio test) to build consensus trees. Finally, phylogenetic trees were visualized using Interactive Tree Of Life (Letunic and Bork 2019) (iTOL, version 4).

### Cluster and enrichment analyses on phylogenetic profile

The phylogenetic profile of human kinases in this study was built upon 558 human kinase domains and their orthologs identified by KinOrtho’s both full-length and domain-based approaches across the 17,134 species in the UniProt reference proteomes (Figure 4). We manually grouped these species into 561 clades based on the NCBI Taxonomy database (Federhen 2012). Each clade contains at least five species; each clade in eukaryotes, bacteria, archaea, or viruses contains at most 41, 240, 35, or 2,287 species, respectively. Then we calculated an ortholog coverage for each kinase-clade pair by dividing the number of orthologs by the total number of species in each clade. Based on this phylogenetic profile (a kinase-clade matrix), in addition to ordering the human kinase domains by their groups defined by KinBase (Figure 4a), we clustered them using k-means clustering (MacQueen 1967) (Figure 4b). We used an R package “factoextra” (Kassambara and Mundt 2017) (version 1.0.7) with options “kmeans” (clustering function), “nstart = 50” (initial random centroids), “nboot = 500” (number of bootstrap samples), and “gap_stat” (compute gap statistic (Tibshirani et al. 2001)) to determine the optimal number of clusters. We found that the optimal number of clusters was 10 (Supplementary Figure S10).

We performed Gene Ontology (GO) enrichment analyses using the GO annotations of all human kinases and their orthologs in each cluster. We extracted all three GO domains (biological process, cellular component, and molecular function) annotations from UniProt (UniProt 2019) (release 2019_11). We then expanded the lowest-level GO terms to all-levels GO terms for every kinase based on the hierarchical controlled vocabulary defined by the GO Consortium (Ashburner et al. 2000; The Gene Ontology 2019). Because the GO terms are annotated at the protein level instead of the domain level, we removed duplicate annotations if a protein’s tandem kinase domains are in the same cluster. When performing enrichment analyses, we chose all kinase orthologs’ annotations as background, used Fisher’s exact test, and then controlled the FDR by the Benjamini-Hochberg procedure (Benjamini and Hochberg 1995). A significantly enriched GO term is defined as its FDR < 0.05, and it is annotated for at least five human kinases in a cluster. If multiple GO terms in the same lineage are enriched in a cluster, we only keep the lowest-level term.

### Novel Inferred Annotation Score

The Novel Inferred Annotation Score (NIAS) proposed in this study is used to estimate the number of potential annotations we can infer from orthologous relationships to annotate understudied human kinases, which are defined by NIH Illuminating the Druggable Genome program (IDG) (Health 2019) (Supplementary Data S4, last updated on June 11, 2019). The scoring system was built upon machine learning-based annotation inference models using overlapping orthologous relationships and GO annotations (Figure 5).

First, to prevent prediction models from being biased by unreviewed data, we only used the manually reviewed (non-electronic) annotations of all well-studied kinases and their orthologs to build training sets for the three GO domains. The GO terms annotated for less than 100 kinases were excluded from the training sets. Each instance of a kinase-GO term pair in the training sets showed the values of output and input features based on the GO annotation status (1 for present and 0 for absent) of a well-studied human kinase and its orthologs, respectively. For example, in Figure 5, an instance shows that mouse’s and rat’s Egfr genes have a GO term “response to stimulus” (GO:0050896) annotation, but human EGFR has not. The training sets consisted of 0.3 million instances with 730 GO terms for 393 well-studied kinases and their orthologs across 176 species. To prioritize the annotations from different species, we further introduced a sequence similarity for each ortholog. Sequence similarities are defined by the average of the normalized weights generated when we built orthologous relationship graphs.

After building the three training sets, we built annotation inference models for each training set using logistic regression, support vector machine (SVM), and random forest implemented by WEKA (Hall et al. 2009). All models were trained with class balancing (using instance reweighting) and 10-fold cross-validation to prevent overfitting. The three machine learning methods’ prediction performances for each training set are shown in Table 2. Because random forest showed the best performance among the three training sets, we used the annotation inference models built by random forest to predict missing GO annotations for understudied human kinases. When building test sets, we used the GO terms annotated for at least one understudied kinase ortholog. There were 11,503 instances in the biological process test set, 1,810 instances in the cellular component test set, and 2,507 instances in the molecular function test set. After applying the random forest models, the confusion matrices built upon the annotation inference result and existing annotations are shown in Supplementary Table S2. We collected those GO annotations currently absent but predicted as present for each understudied kinase and then only kept the lowest-level term in each GO term lineage. The summation of each prediction score calculated by random forest is defined as an Inferred Annotation Score (IAS):

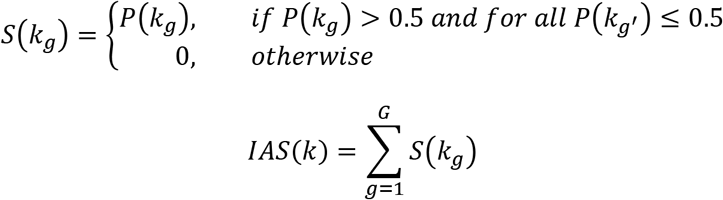

where *P*(*k_g_*) is the prediction score of *g*th GO term annotation for understudied kinase *k, g* = {1,2,…, *G*}, and *g′* represents any descendant of *g*th GO term. Because the predicted annotations may include existing unreviewed electronic annotations, we defined the NIAS of an understudied kinase by subtracting the unreviewed annotation score (UAS) from IAS:

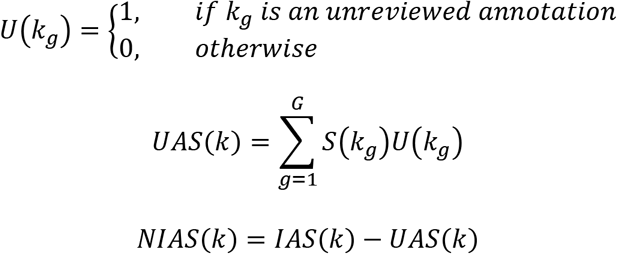

## Data access

All source codes are available at https://github.com/esbgkannan/KinOrtho

## Acknowledgements

This work was supported by National Institutes of Health (funding for NK from U01CA239106). NG was supported by the NIGMS under award number R25GM109435. The funding bodies did not play any roles in the design of the study and collection, analysis, and interpretation of data and in writing the manuscript. We would like to thank Dr. Karen Ross, members of the IDG consortium and NK Lab for useful comments on the manuscript.

## Author contributions

LH, RT, and NK designed the research. LH and NG built the software. LH and RT evaluated the software’s performance. LH and RT performed phylogenetic analysis. LH performed enrichment analyses. LH performed machine learning methods. LH, RT, WY, and NK analyzed the data and interpreted the results. LH, RT, and AV wrote the manuscript. LH created the tables and figures. LH, NG, WY, DPB, PAE, and NK revised the manuscript. All authors reviewed the manuscript and approved the final version of the manuscript.

